# Evaluating a digital sepsis alert in a London multi-site hospital network: a natural experiment using electronic health record data

**DOI:** 10.1101/637967

**Authors:** Kate Honeyford, Graham S Cooke, Anne Kinderlerer, Elizabeth Williamson, Mark Gilchrist, Alison Holmes, The Sepsis Big Room, Ben Glampson, Abdulbrahim Mulla, Ceire Costelloe

**Author notes:** **Corresponding author:**, Global Digital Health Unit, Department of Primary Care and Public Health Imperial College, Charing Cross Campus, Reynolds Building, St Dunstan’s Road, London, W6 8RP, UK.

## Abstract

**Objective:** To determine the impact of a digital sepsis alert on patient outcomes in a UK multi-site hospital network.

**Methods:** A natural experiment utlising the phased introduction of a digital sepsis alert into a multi-site hospital network. Sepsis alerts were either visible to clinicans (the ‘intervention’ group) or running silently and not visible (the control group). Inverse probability of treatment weighted multivariable logistic regression was used to estimate the effect of the intervention on patient outcomes.

Outcomes: In-hospital 30-day mortality (all inpatients), prolonged hospital stay (≥7 days) and timely antibiotics (≤60 minutes of the alert) for patients who alerted in the Emergency Department.

**Results:** The introduction of the alert was associated with lower odds of death (OR:0.76; 95%CI:(0.70, 0.84) n=21,183); lower odds of prolonged hospital stay ≥7 days (OR:0.93; 95%CI:(0.88, 0.99) n=9988); and in patients who required antibiotics, an increased odds of receiving timely antibiotics (OR:1.71; 95%CI:(1.57,1.87) n=4622).

**Discussion:** Current evidence that digital sepsis alerts are effective is mixed. In this large UK study a digital sepsis alert has been shown to be associated with improved outcomes, including timely antibiotics, which may suggest a causal pathway. It is not known whether the presence of alerting is responsible for improved outcomes, or whether the alert acted as a useful driver for quality improvement initiatives.

**Conclusions:** These findings strongly suggest that the the introduction of a network-wide digital sepsis alert is associated with improvements in patient outcomes, demonstrating that digital based interventions can be successfully introduced and readily evaluated.

**Funding:** Imperial NIHR Biomedical Research Centre: NIHR-BRC-P68711.

## BACKGROUND

Sepsis is recognised as a common cause of serious illness and death. It is estimated that there are 123,000 cases in England each year and 46,000 deaths.[1] Similar high levels of sepsis have been reported internationally [2–3] and sepsis is recognised by WHO as a global health priority.[4] Many countries have nationwide sepsis action plans and in England there are targets for hospitals to rapidly diagnose and treat patients with sepsis.

Timely, appropriately targeted, intravenous (IV) antibiotics have been shown to be effective in improving outcomes for patients, with a 4% increase in odds of mortality for every hour’s delay in administration of IV-antibiotics.[5–7] This evidence has resulted in UK hospitals having a target (with financial incentives) of sepsis patients receiving IV antibiotics in one hour.[8–9]

In order to ensure rapid diagnosis and early treatment sepsis screening tools have been introduced and refined, these include qSOFA,[10] NEWS, [11] and NEWS2.[12] Early Warning Scores (EWS) have been shown to be effective in predicting mortality [13] and ICU admission.[14] There is limited evidence that the introduction of track and trigger style warning systems have been associated with improved outcomes for patients.

The introduction of electronic health records (EHRs) has provided the opportunity to embed digital alerts based on current and past clinical measurements. A range of screening algorithms have been used, including the St John Sepsis Algorithm (SJSA) [15–16], the Severe-Sepsis Best Practice Alert [17] and hospital designed alerts.[18] The evidence for the effectiveness of these alerts on patient outcomes is mixed. Some studies have shown that introduction of digital sepsis alerts have led to increases in the proportion of patients with suspected sepsis receiving IV-antibiotics in one hour,[17] reduced ICU and hospital length of stay [19] and reduced in-hospital mortality,[18–19] whilst others have shown no significant effect on length of stay [19–20] or in-hospital mortality.[17] A recent randomized control trial (RCT) analysing the impact of the introduction of an alert for inpatients in a US hospital found no association between the introduction of an alert and an improvement in patient outcomes, although the study was underpowered.[21] The majority of evidence comes from small, ICU based studies in the US healthcare system. It is not known if similar impacts on patient outcomes will be seen in larger scale studies, particularly in an English hospital, where care is free at the point of delivery and accessible to all.

## OBJECTIVE

The aims of this study were to determine the effect of the introduction of a digital sepsis alert on one key process measure (timely antibiotics) and two patient outcomes (extended length of stay and in-patient 30-day mortality).

## METHODS

### Study design

In this natural experiment, a weighted multiple logistic regression was used to examine the effect of the digital sepsis alert. Data from October 2016 to May 2018 was included in the study which utilized a ‘silent’ running phase, during which time digital alerts were active, but were not visible to clinicians. The silent phase provides a control group. Robust statistical methods were used to balance characteristics between the live and control phases. The primary outcome was 30-day inpatient mortality and a secondary outcome of prolonged length of stay (≥7 days). Additionally, the impact of the introduction of the alert on the key process measure of timely IV-antibiotics (≤60 minutes after the alert) was studied.

### The digital sepsis alert

The digital sepsis alert is based on the St John Sepsis Algorithm developed by Cerner,[24] shown in Figure 1. The alert is an integrated part of the Electronic Health Record (EHR) and has a silent running mode. Silent alerts are not visible to clinical staff at the ‘front end’ of the system. Once the alert is turned on (live) in a clinical area nurses and doctors are notified of patients who have triggered the alert. Nurses are notified of patients who have triggered the alert either through a pop-up warning on the EHR (in inpatient wards) and/or as a dashboard which highlights any patient with an active alert (in the ED and inpatient wards). Doctors are presented with a sepsis warning only when they open the patient’s record. In addition to the alert, a novel multidisciplinary-care-pathway, designed by the Trust, launches from the digital record when the clinician confirms suspicion of sepsis. These ‘Treatment Plans’ support the clinician to start treatment in-line with hospital guidance, including fluids, oxygen, diagnostic tests (blood and other cultures), and early antibiotics. Content from local infection guidelines is built in, so that for any given potential sepsis diagnosis, the appropriate antibiotics with appropriate dosing, and directions are prompted.

**Figure 1.**
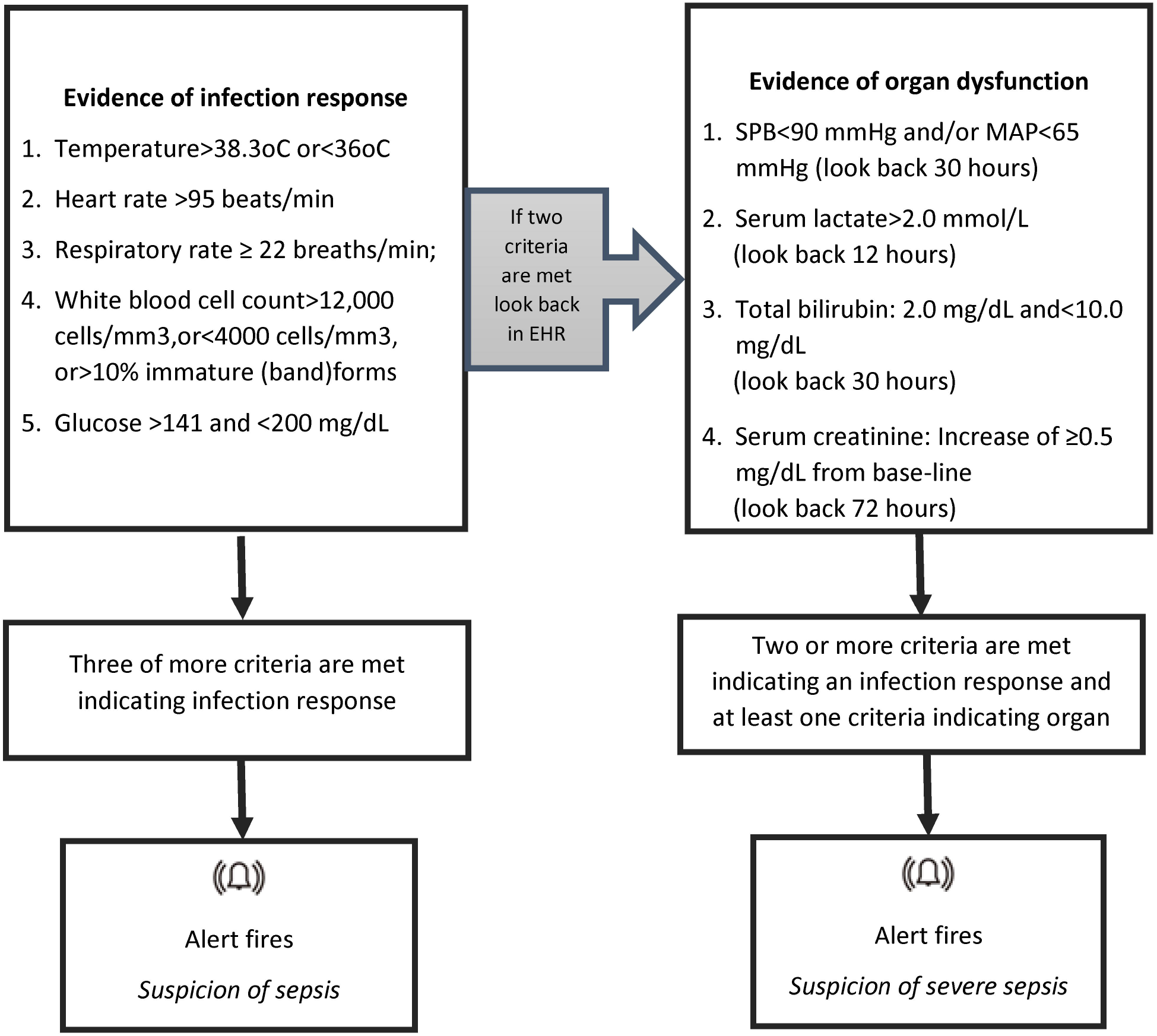
Criteria associated with the St John’s Sepsis Alert [23]

The introduction of the alerts was part of a framework for system redesign and improvement, see Box 1 for more details. The alert was introduced in a phased approach over an 18 month period across the Trust, summarised in Figure 2. The alert was switched from silent to live as recommended by improvement approaches for scale and spread.[25] Initially in the acute medical unit at one site, expanding out to both Emergency Departments and haematology wards and then Trust-wide. Initial areas were selected based on their interest in assistance in identifying patients with suspicion of sepsis.

**Figure 2.**
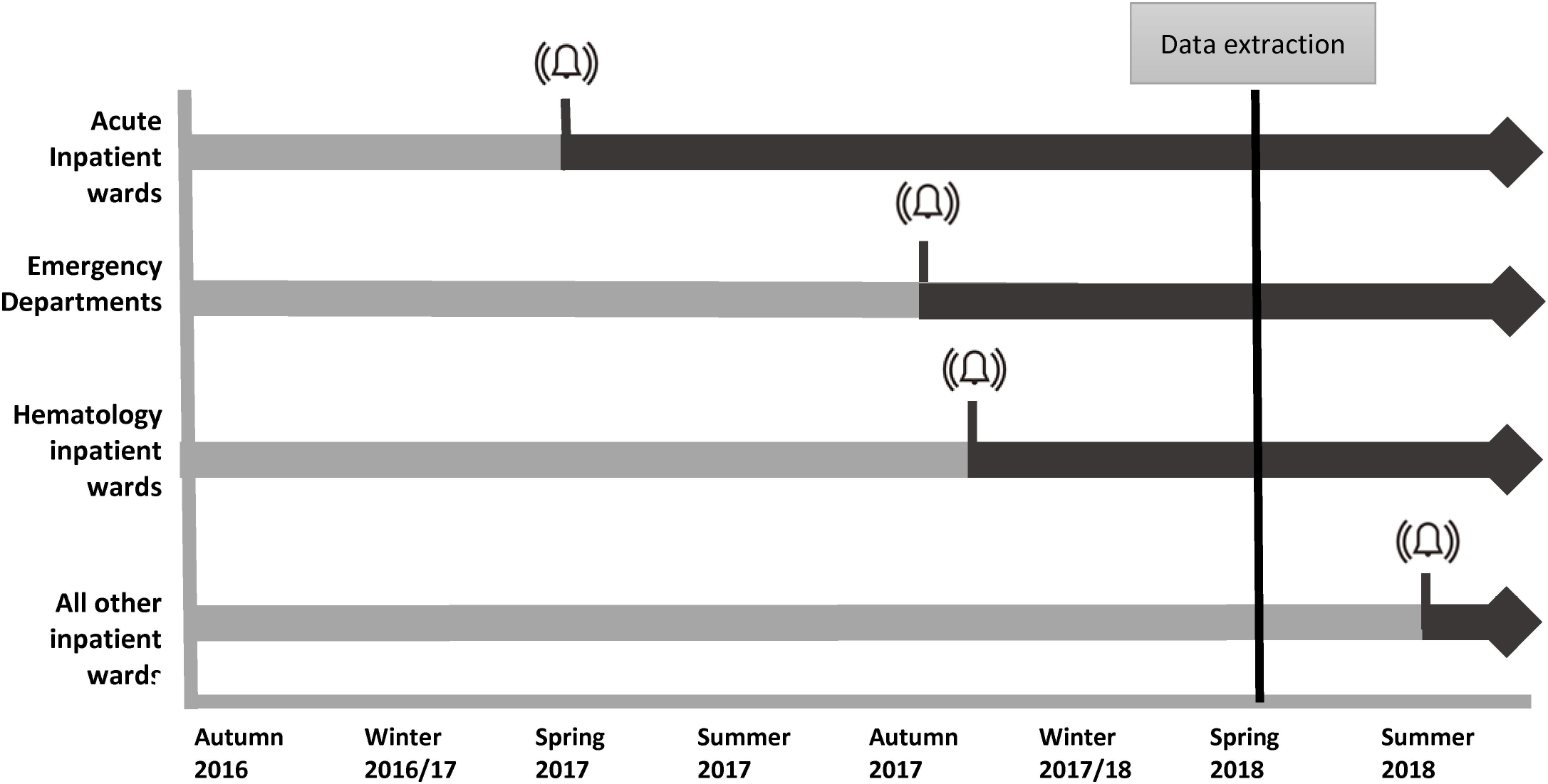
Phased introduction of live alerts across multi-site hospital from Autumn 2016. The digital alert was switched from silent to live in acute wards, followed by Emergency departments in two hospital sites in Autumn and haematology departments soon after. The alert was switched to love across all inpatient wards in August 2018, after data was extracted for this study.

#### Box 1 The local picture

The hospital network (Trust) comprises of five main sites. In recent years, the Trust had over one million outpatient contacts, quarter of a million A&E attendances, 200,000 inpatient contacts and 100,000 inpatient operations. The Trust employs over 2500 doctors, 4,400 nurses, 720 allied healthcare professionals and 130 pharmacists.

Work to improve care for sepsis patients at the Trust centres on three key priorities:

- The identification and treatment of sepsis across the whole patient pathway
- Consistency of standards and reporting
- The prudent use of antimicrobials within the wider antimicrobial stewardship and resistance agenda.

A key focus has been to ensure patients identified with sepsis receive the appropriate antibiotics within one hour, in line with national targets. The work is integrated with the digital transformation and the use of an embedded digital sepsis alert in the EHR. The digital alert was initially embedded in the EHR running ‘silently’, that is, not visible to clinical staff. An analysis was planned to evaluate the impact of making the alert visible to clinical staff. Implementation of the alert was part of a collaborative improvement approach through the Sepsis Big Room. A “big room” is a weekly coached meeting which provides time and space for a range of staff to come together to discuss improvements to the quality of patient care. Staff from all disciplines are welcome and the meetings operate a flattened hierarchy. Patient stories are reviewed and real-time data displayed to support the identification of specific improvements to healthcare processes within the pathway of care. In an approach similar to that others have used, a series of tests of change were undertaken to improve decision making and communication for sepsis patients. For each test, a small-scale Plan Do Study Act (PDSA) cycle, based on Toyota Big Room methodology,[22–23] was performed and, if this proved successful, the test was tried more widely. Data from the evaluation, was used to provide feedback to the team and shape discussion on implementation.

### Patient population

Patients who triggered the alert were included in the analysis. These are patients who are identified as potentially having sepsis by the clinical thresholds included in the St John’s Sepsis alert (Fig 1). The unit of analysis is an adult inpatient ‘encounter’. An encounter was defined as a continuous spell in the Trust. In this analysis all encounters of adult patients (aged 18 an over) in which a sepsis alert was triggered were included. Although the alert may be triggered repeatedly for a patient during a hospital encounter, only first alerts were considered. All patients admitted to the three hospitals in the network that have general acute admissions were included. Only encounters ending in admission were included in final models to ensure detailed comorbidity and outcome information were available. For length of stay and timely antibiotics only patients who alerted in the ED were included, this is to reduce potential confounders such as the use of prophylactic antibiotics and the impact of prior long inpatient stays on LOS post infection. Patients who were already on antibiotics were excluded from the analysis of timely antibiotics. Patients who did not receive ABX within 24 hours of the alert were excluded it was assumed the alert had triggered for patients who did not require antibiotics (further details in Supplementary Materials S1). See Figure 3 for a summary of patient cohorts.

**Figure 3.**
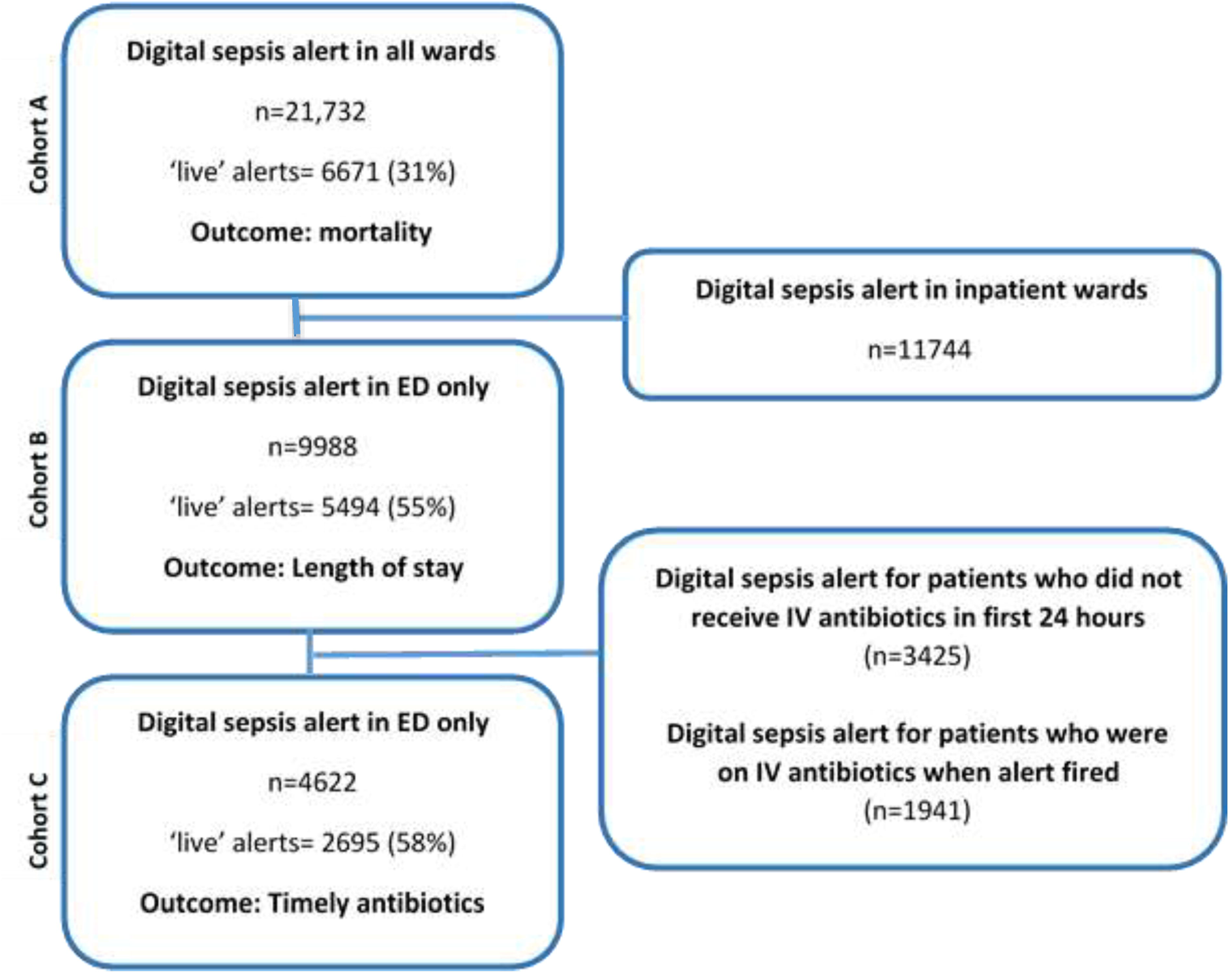
Cohort definition. Three cohorts were developed to investigate the outcomes of interest: Cohort A comprised of all patients who alerted and the outcome of interest in this cohort was mortality. Cohort B comprised of patients who alerted in the emergency departments only as the main outcome of interest was length of stay. Cohort C comprised of patients who alerted in the emergency department who received antibiotics within 24 hours post alert. The main outcome of interest was – timely antibiotics, defined a receiving antibiotic within 1 hour of the alert (as per NICE guidelines).[25]

### Outcomes

The outcomes were: (i) in-hospital all-cause mortality within 30 days of alert (ii) long hospital stay (≥ 7 days) and (iii) timely antibiotics (IV-antibiotics ≤60 minutes). Both long hospital stay and timely antibiotics were investigated only for patients who alerted in the ED. Length of stay was measured as time from alert to discharge. For the purposes of this study ‘timely antibiotics’ was defined as patients who received IV-antibiotics within one-hour of the alert. This definition was informed by the current target for hospitals in England.[8]

### Statistical analysis

Three separate analyses were undertaken in three cohorts to explore the three main outcomes. The switch from silent to live was considered as a natural experiment. Inverse probability of treatment weighting (IPTW) was used [27] to account for confounding the non-random allocation introduced and balance characteristics between the live and control phases. Multivariable logistic regression was used to determine the propensity score weights. Further details are available in Supplementary Materials (S2). Potential confounders for the three outcomes and alert status allocation were included in models. These included patient characteristics, admission details and clinical measures. Data were obtained from patient digital medical records. Details of variables are included in Box 2.

#### Box 2 Covariates included in both the propensity score models and the multivariable logistic models.

| Patient characteristics |  |
| --- | --- |
| **Age** | Grouped into 18-44; 45-64; 65-69; 70-74; 75-79; 80-84; and 85+ |
| **Sex** | Male or female |
| **Ethnicity** | Based on the following groupings:<br>White; Asian; Black; other; and not known.<br>*More detail to be added here.* |
| **Comorbidities** | Based on any relevant ICD-10 code appearing in the discharge diagnosis codes (25 possible codes)<br>MI; CHF; PVD; Stroke; Dementia; Pulmonary; Rheumatic; PUD; Liver (mild); Liver (severe); Diabetes; Diabetes (complex); Paralysis; Renal; Metastatic cancer; HIV. |
| **Deprivation quintile** | Measured as the deprivation score of the patient's GP practice, obtained by matching patients to their registered GP. If patients did not have a registered GP or the practice was not included in the PHE practice profiles a 'missing' categorisation was allocated. There are therefore six deprivation categories, with Quintile 1 being the least deprived. |
| Admission characteristics |  |
| **Admitting hospital** | A has an A&E department<br>B no A&E department<br>C has an A&E department |
| **Season of admission** | Spring: March, April, May<br>Summer: June, July, August<br>Autumn: September, October, November,<br>Winter: December, January, February |
| Patient Severity |  |
| **NEWS SCORE** | Categorised into:<br>Zero, Low, Medium, High and none recorded<br>NEWS = 0 : Zero<br>1 ≤ NEWS < 5 : Low<br>5 ≤ NEWS < 7 : Medium<br>7 ≥ NEWS : High<br>A NEWS score is available for 19599 of the patients (90% of the patients). |
| **Alert** | Suspected sepsis<br>Suspected severe sepsis |

Balance between treatment populations was evaluated using standardised mean differences (SMDs) of all baseline covariates. A threshold of 10% indicates possible imbalance, and 25% as an indication of unacceptable imbalance.

Odds ratios and 95% confidence intervals were estimated for each outcome using logistic regression applying the propensity score weights. A doubly robust approach was employed,[28] including covariates in both the propensity score models and the multivariable logistic models of the outcomes. When modelling death, a random-effects model was used to account for clustering within the acute, hematology and ED areas, with all other alerts included in an additional cluster. Additionally the outcomes were modeled using logistic regression without applying the propensity score weights but adjusting for confounders. All analyses were done with R V.3.2.3 (R Foundation for Statistical Computing, Vienna, Austria).

## RESULTS

### Study Population

In total there were 21,732 patient encounters with at least one alert between October 2016 and May 2018. 9988 of these were in the ED, 942 alerted in acute wards and 1218 in haematology wards. 4622 ED patients were not on IV-antibiotics at the time of the alert and did receive IV-antibiotics within 24 hours of the alert. See Figure 3 for cohort details.

Table 1 summarises the characteristics of patients for whom the alert was during a silent phase and those for whom the alert was in a live phase – clearly visible to clinicians. The phased introduction of the live alert means the patient and encounter characteristics of the two groups are not balanced and there are more live alerts for patients admitted to Site C, admitted through the ED and admitted in the autumn and winter. Within live alerts there was a higher proportion of younger patients (18-44) and older patients (85 and over). In comparison to silent running, patients who alerted in the live phase were more likely to have pulmonary conditions, but less likely to have renal conditions. In addition they were less likely have an unknown ethnicity. A higher proportion of patients who alerted in the live phase had medium, high or missing NEWS score. A higher proportion had suspected severe sepsis in comparison to suspected sepsis.

**Table 1.**
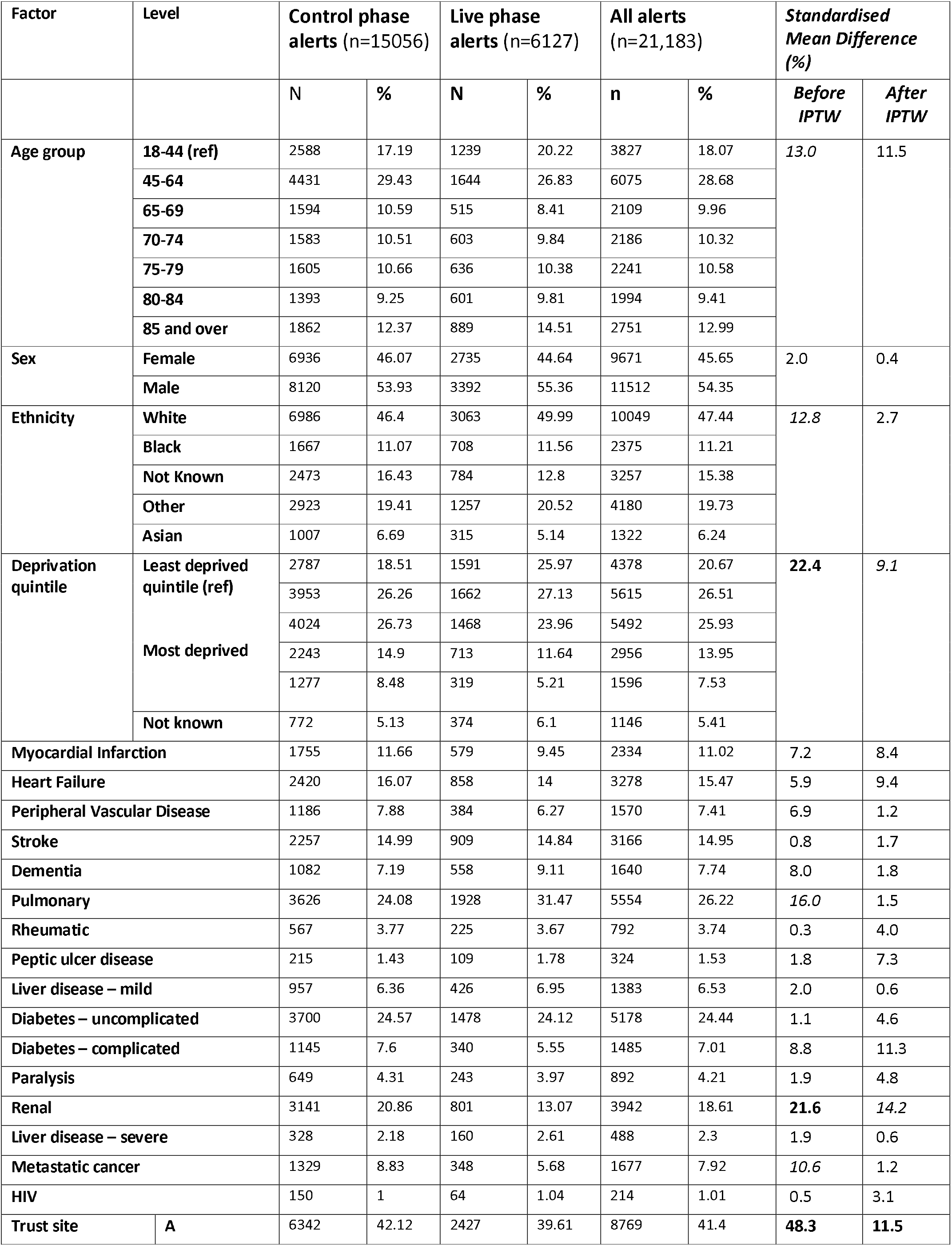
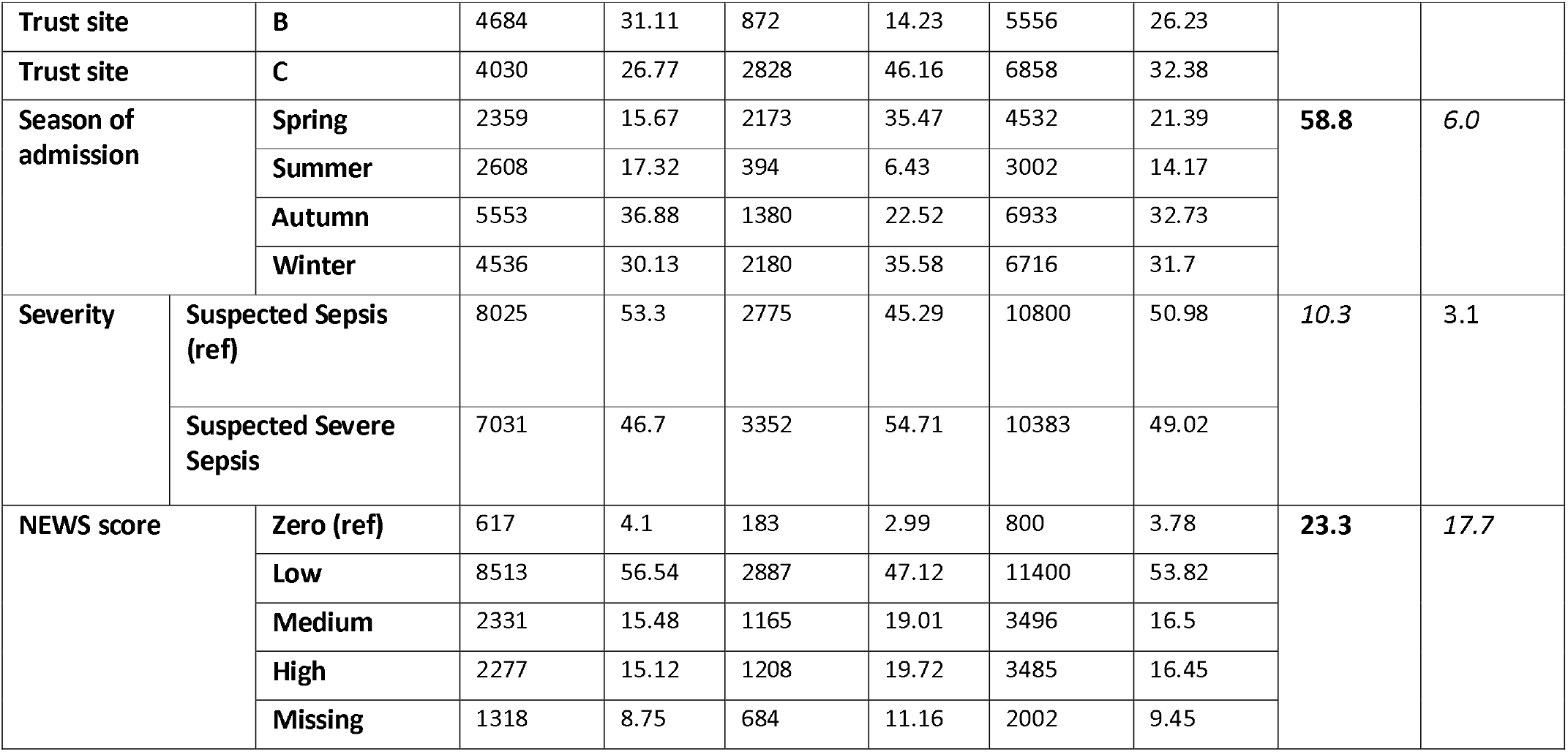
Distribution of patient and encounter characteristics for all alerts and standardised mean difference before and after weighting.

Inverse probability of treatment weight was used to account for these baseline differences. Models were adjusted for differences in the two groups using IPTW. After weighting all models were balanced. SMDs for cohort B used to model length of stay and cohort C used to model timely antibiotics were 3.0% or less (compared with a highest value of over 70% before imposing estimated weights). For cohort A, to model death, the SMDs for hospital site, alert introduction area (cluster), age, NEWS, diabetes and renal disease, were all above 10% but under 25%. As a doubly robust method all potential confounders were included in the final model accounting for any confounding from known confounders. Further details are available in Supplementary Materials (S3).

### Association of alert status with death

21732 inpatients alerted during the period of study, across all wards, across the three sites. 1293 (6%) patients died within 30 days of the alert being triggered, which is similar to mortality rates reported elsewhere. [21] Live alert status was associated with lower in-hospital mortality (5.1% compared to 6.4%, p<0.001), see Table 2. After accounting for patient characteristics, using IPTW propensity scores and patient characteristics in the multivariable logistic model, patients who alerted during the live phase had 24% lower odds (95% confidence interval (CI): 16 to 30% lower odds) of in-hospital death.

**Table 2.** Summary data and results of models, including adjustment for confounders (adjusted (reg)^1^). After adjustment for potential confounding and IPTW measures^2^ of association did not change markedly, but were more precise

|  | Death |  | Extended LOS |  | Timely ABX |  |
| --- | --- | --- | --- | --- | --- | --- |
|  | Control | Live | Control | Live | Control | Live |
| **Total encounters** | 15061 | 6671 | 4494 | 5494 | 1927 | 2695 |
| **Number events** | 959 | 339 | 1846 | 2209 | 712 | 1204 |
| **% events** | 6.4 | 5.1 | 41.1 | 40.2 | 36.9 | 44.7 |
|  | **OR (95% CI)** |  | **OR (95% CI)** |  | **OR (95% CI)** |  |
| **Unadjusted** | 0.67 (0.67, 0.90) |  | 0.97 (0.89, 1.05) |  | 1.38 (1.22, 1.55) |  |
| **Adjusted (reg)<sup>1</sup>** | 0.79 (0.67, 0.93) |  | 0.97 (0.87, 1.05) |  | 1.70 (1.43, 1.95) |  |
| **Adjusted (IPTW)<sup>2</sup>** | 0.76 (0.70, 0.84) |  | 0.93 (0.88, 0.99) |  | 1.71 (1.57, 1.87) |  |
|  | **RR<sup>3</sup> (95% CI)** |  | **RR<sup>3</sup> (95% CI)** |  | **RR<sup>3</sup> (95% CI)** |  |
| **Adjusted (IPTW)<sup>2</sup>** | 0.76 (0.70, 0.84)<sup>3a</sup> |  | 0.96 (0.93, 0.99) |  | 1.35 (1.28 to 1.41) |  |

<sup>1</sup>Adjusted for all confounders summarised in Table 1.
<sup>2</sup>Propensity score weighted log linear fully adjusted model used to estimate relative risks.
<sup>3</sup>Relative risks determined using a fully adjusted, propensity score weighted log-linear model. <sup>3a</sup>With the exception of death which is estimated directly from the odds ratio as the event is rare.

### Association of alert status with long length of stay (LOS)

9988 patients alerted in the ED and were subsequently admitted and 4055 (40.6%) of these subsequently had a long length of stay (≥ 7 days), measured as time from the alert to discharge. After accounting for patient characteristics, live alert status was significantly associated with decreased odds of long length of stay. Patients who alerted during the live phase had a 7% lower odds of a long length of stay.

### Association of alert status with timely antibiotics

6563 of the 9988 (65.7%) patients who alerted in the ED received antibiotics within plus or minus 24 hours of the alert firing.

Of the 4622 patients who were not on antibiotics when the alert fired, 36.9% of encounters which activated during the control period resulted in IV-antibiotics administered within one hour of the alert, and 44.7% of encounters that activated the alert when it was visible to clinicians. After accounting for patient characteristics, live alert status was associated with 71% higher odds of receiving timely antibiotics (odds ratio (OR):1.71; 95%CI:1.57 to 1.87). This approximates to a relative risk of 1.35 (95%CI: 1.28 to 1.41); a 35% increase in chance of receiving timely IV-antibiotics.

Patients who did not receive antibiotics are summarised in Table S1. A in the Supplementary Materials. It is assumed that the majority of these patients did not need IV-antibiotics. This assumption is supported by the improved prognosis of these patients (6.4% died and 46.6% had a prolonged LOS compared to 3.8% and 29.1% of those who did not receive antibiotics). Patient characteristics were compared in those that received antibiotics and those that did not. A higher proportion of elderly patients, patient with a high NEWS score, patients who alerted in the spring and winter received antibiotics. Ethnicity, deprivation and sex were not significantly associated with receipt of antibiotics, suggesting that clinical aspects of the patient and not underlying health inequalities are associated with receipt of antibiotics.

## DISCUSSION

This is the first evaluation of a digital sepsis alert in an English hospital and the largest undertaken anywhere to date. Robust methods were used to show that the introduction of a digital sepsis alert was associated with improvements in patient outcomes. Overall, 6% of patients who alerted as possible sepsis patients died within 30 days of the alert (all-cause, in-hospital), this is in a hospital network with a lower than expected overall in-hospital mortality.[29] Patients for whom their first alert was during the live phase had lower odds of death (OR: 0.76; 95% CI:(0.70, 0.84)), lower odds of a long hospital stay (≥7 days) (OR: 0.93; 95% CI:(0.88, 0.99)) and increased odds of receiving timely antibiotics (OR:1.71; 95%CI:(1.57, 1.87)). The magnitude and interpretation of these results is similar when using either a weighted or unweighted multiple logistic regression model. These results suggest an important clinical benefit from the introduction of alerting, although it is not possible to say the extent to which the presence of alerting, per se, is responsible for the benefits seen, or whether the alert acted as a useful driver for other quality improvement initiatives.

The phased nature of the introduction of the alert allows a detailed analysis, with live alerts and controls coming from a range of wards, settings and time periods. Smaller studies in the US have found interventions which included an alert triggered by an EHR based diagnosis resulted in improved outcomes for patients. McRee et al [20] found a reduction in mortality from 9.3% to 1.0%, but no significant effect on length of stay, although this was a small, pilot study (n=171). In another small study (n=214) in the US the introduction of an alert was associated with an improvement in patients receiving timely antibiotics, from 48.6% to 76.7%, and a significant reduction in length of stay. [17] There was no significant impact on in-hospital mortality, which may be explained by the low numbers in the study. Westra [30] and Guirgis [31] found the introduction of an alert, bundled with education, training and structured care sets, to be associated with reductions in mortality and length of stay. Manaktala [18] report a 53% reduction in mortality, although no significant impact on length of stay, when an alert was introduced into specific units which had received training. In addition to observational studies, a recent RCT found the introduction of a sepsis alert had no impact on receipt of antibiotics within three hours. However, this study only included patients in wards, not the ED, and as the RCT was terminated early there was insufficient statistical power to detect change.[21]

This analysis of a large sample of patients who have been admitted across three sites to a busy hospital network in England is one of the largest to date. For mortality both patients who presented with suspected sepsis in the ED and those who developed symptoms congruent with sepsis during their inpatient stay were included. For length of stay and timely antibiotics patients who alerted in the ED and were subsequently admitted were included in the analysis. Outcomes were selected based on their importance to both patients and hospitals, including those based on UK government targets, and were applied to all patient encounters where there was an alert, not limited to patients who were confirmed as having sepsis. A key methodological strength of this study was the inclusion of a ‘silent’ running phase, during which time digital alerts were active, but were not visible to clinicians. The silent phase provides a control group. In addition, robust statistical methods were used to balance characteristics between the live and control phases.

There are a number of limitations to our study. Firstly, the quasi-experimental design limits ability to imply causation. Although RCTs are considered the gold standard for analyzing health interventions this approach was not deemed possible in the complex environment of this busy, multi-site hospital. Difficulties of conducting a RCT on digital alerts for sepsis have been documented elsewhere.[21] Statistical approaches recommended for the analysis of data derived from natural experiment were used.[32] Propensity scores are a recognized and recommended method to adjust for confounding factors, in this case introduced by the phased introduction of the alert. The majority of live alerts were ED patients who attended in autumn and winter which resulted in a higher proportion of severe sepsis and higher NEWS scores in the live group. The impact of the wider sepsis Quality Improvement (QI) initiatives on improved outcomes for patients could not be robustly modelled as it was not possible to associate the introduction of these initiatives to specific periods of time. Aspects of our analysis were limited by data availability; only admitted patients were included because clinical information was limited for patients discharged from the ED without admission. The analysis of the association of the alert on timely antibiotics and LOS was limited to patients in the ED to reduce potential confounders such as the use of prophylactic antibiotics and the impact of prior long inpatient stays on LOS post infection.

In this study we have not considered unintended consequences of the introduction of the alert, including the possibility of increases in the use of inappropriate and/or unnecessary IV antibiotics, increases in patients being diagnosed as having sepsis without confirmation, and the possibility of alert fatigue as a result of relatively low specificity of the alert. This is the focus of future work.

## SIGNIFICANCE

The introduction of digital altering for sepsis is an opportunity to improve care for patients who may have sepsis. Despite the current emphasis on the use of sepsis screening tools, including the recommendation of the uptake of NEWS2 as the best current option for standardising the management of deterioration and sepsis,[33] there is uncertainty around the use of digital screening to improve patient outcomes.[34] This study, the largest to date is an important addition to the body of knowledge that appropriate digital based screening, when associated with quality improvement approaches is associated with improved patient outcomes.

## CONCLUIONS

In two busy acute hospitals, the introduction of a digital sepsis alert has been shown to be associated with improved patient outcomes, including lower risk of mortality and extended length of stay. A 70% increase in odds of receiving timely antibiotics was found, which is likely to be important in explaining the causal pathway for the alert improving outcomes for patients. This study has clearly shown that the introduction of a network-wide digital screening tool embedded in EHRs is associated with improvement in patient outcomes, demonstrating that digital based interventions can be successfully introduced and readily evaluated.

## Supporting information

who did not require antibiotics (further details in Supplementary Materials S1

## ADDITIONAL INFORMATION

### Acknowledgements

We thank members of Imperial College NHS Healthcare Trust, specifically the Sepsis Big Room, who participated in the study. We acknowledge the National Institute of Health Research Imperial Biomedical Research Centre who funded this study.

The research was conducted using NIHR HIC data resources.

This work was carried out in collaboration with the National Institute for Health Research Health Protection Research Unit (NIHR-HPRU) in Healthcare Associated Infection and Antimicrobial Resistance at Imperial College London.

### Disclaimer

The views expressed are those of the author(s) and not necessarily those of the NHS, the NIHR or the Department of Health and Social Care.

### Author Contributions

CC & GC conceived the study. KH, CC, GC & AK developed the protocol. AM, BG, MG, AK, CC and KH defined the variables of interest from the clinical data and extracted the data. KH conducted the statistical analysis which was reviewed and improved by EW and CC. MG and AK, working with the Big Room refined and developed the data definitions and their clinical relevance and provided ongoing feedback on results and their interpretation. KH wrote the first draft of the manuscript. All authors reviewed and contributed to the final draft of the manuscript. CC is the guarantor. The corresponding author attests that all listed authors meet authorship criteria and that no others meeting the criteria have been omitted.

## BIBLIOGRAPHY

1 NHS England. Overview Sepsis 2016. https://www.nhs.uk/conditions/sepsis/. Accessed 28 December 2018.

2 Paoli CJ, Reynolds MA, Sinha M, Gitlin M, Crouser E. Epidemiology and Costs of Sepsis in the United States-An Analysis Based on Timing of Diagnosis and Severity Level. Crit Care Med. 2018.

3 Prescott HC, The Epidemiology of Sepsis, in Wiersinga W & Seymour C (ed) Handbook of Sepsis 2018

4 Reinhart K, Daniels R, Kissoon N, Machado FR, Schachter RD, Finfer S. Recognizing Sepsis as a Global Health Priority - A WHO Resolution. The New England Journal of Medicine. 2017;377(5):414–7.

5 Kumar A, Roberts D, Wood KE, Light B, Parrillo JE, Sharma S, et al. Duration of hypotension before initiation of effective antimicrobial therapy is the critical determinant of survival in human septic shock. Crit Care Med. 2006;34(6): 1589–96.

6 Ferrer R, Martin-Loeches I, Phillips G, Osborn TM, Townsend S, Dellinger RP, et al. Empiric antibiotic treatment reduces mortality in severe sepsis and septic shock from the first hour: results from a guideline-based performance improvement program. Crit Care Med. 2014;42(8):1749–55.

7 Seymour CW, Gesten F, Prescott HC, Friedrich ME, Iwashyna TJ, Phillips GS, et al. Time to Treatment and Mortality during Mandated Emergency Care for Sepsis. The New England Journal of Medicine. 2017;376(23):2235–44.

8 NHS CQUIN Indicator Specification Information on CQUIN 2017/18 - 2018/19 2019 Gateway Reference 07725 https://www.england.nhs.uk/wp-content/uploads/2017/07/cquin-indicator-specification-information-january-2019.pdf Accessed: 29 April 2019

9 lacobucci G. NHS hospitals could be fined if they miss new sepsis targets BMJ 2019; 364: l1124

10 Singer M, Deutschman CS, Seymour CW, et al. The Third International Consensus Definitions for Sepsis and Septic Shock (Sepsis-3). JAMA. 2016;315(8):801–810. doi:10.1001/jama.2016.0287Singer, Deutschman, Seymour, et al., JAMA, 2016

11 Royal College of Physicians. National Early Warning Score (NEWS) Standardising the assessment of acute-illness severity in the NHS. London: RCP; 2012.

12 Royal College of Physicians. National Early Warning Score (NEWS) 2: Standardising the assessment of acute-illness severity in the NHS. Updated report of a working party. London: RCP; 2017.

13 Gupta T, Puskarich MA, DeVos E, Javed A, Smotherman C, Sterling SA, et al. Sequential Organ Failure Assessment Component Score Prediction of In-hospital Mortality From Sepsis. J Intensive Care Med. 2018:885066618795400.

14 Goulden R, Hoyle MC, Monis J, Railton D, Riley V, Martin P, et al. qSOFA, SIRS and NEWS for predicting inhospital mortality and ICU admission in emergency admissions treated as sepsis. Emerg Med J. 2018;35(6):345–9.

15 Amland RC, Sutariya BB. Quick Sequential [Sepsis-Related] Organ Failure Assessment (qSOFA) and St. John Sepsis Surveillance Agent to Detect Patients at Risk of Sepsis: An Observational Cohort Study. Am J Med Qual. 2018;33(1):50–7.

16 Amland RC, Sutariya BB. Saving lives through sepsis surveillance. Cerner. 2017. https://www.cerner.com/blog/saving-lives-through-sepsis-surveillance/. Accessed 28 December 2018.

17 Narayanan N, Gross AK, Pintens M, Fee C, MacDougall C. Effect of an electronic medical record alert for severe sepsis among ED patients. Am J Emerg Med. 2016;34(2):185–8.

18 Manaktala S and R Claypool SR, Evaluating the impact of a computerized surveillance algorithm and decision support system on sepsis mortality, Journal of the American Medical Informatics Association, Volume 24, Issue 1, January 2017, Pages 88–95, https://doi.org/10.1093/jamia/ocw056

19 Guirgis FW, Puskarich MA, Smotherman C, Sterling SA, Gautam S, Moore FA, et al. Development of a Simple Sequential Organ Failure Assessment Score for Risk Assessment of Emergency Department Patients With Sepsis. J Intensive Care Med. 2017:885066617741284.

20 McRee L, Thanavaro JL, Moore K, Goldsmith M, Pasvogel A. The impact of an electronic medical record surveillance program on outcomes for patients with sepsis. Heart Lung. 2014;43(6):546–9.

21 Downing NL, Rolnick J, Poole SF, et al Electronic health record-based clinical decision support alert for severe sepsis: a randomised evaluation BMJ Qual Saf Published Online First: 14 March 2019. doi: 10.1136/bmjqs-2018-008765

22 Silvester KM, Mohammed MA, Harriman P, Girolami A, Downes TW. Timely care for frail older people referred to hospital improves efficiency and reduces mortality without the need for extra resources. Age Ageing. 2014;43(4):472–7.

23 Offord N, Harriman P, Downes T. Discharge to assess: transforming the discharge process of frail older patients. Future Healthcare Journal. 2017;4(1):30–2.

24 Amland RC, Hahn-Cover KE. Clinical Decision Support for Early Recognition of Sepsis. Am J Med Qual. 2016;31(2):103–10.

25 Greenhaigh T, Papoutsi C. Spreading and scaling up innovation and improvement BMJ 2019; 365: l2068

26 NICE National Institute for Clinical Excellence. Sepsis Quality standard [QS161]: NICE; 2017. https://www.nice.org.uk/guidance/qs161. Accessed 28 December 2018.

27 Austin PC, Stuart EA. Moving towards best practice when using inverse probability of treatment weighting (IPTW) using the propensity score to estimate causal treatment effects in observational studies. Stat Med. 2015;34(28):3661–79.

28 Lunceford JK. Stratification and weighting via the propensity score in estimation of causal treatment effects: a comparative study. Stat Med. 2017;36(14):2320.

29 NHS Digital. Summary Hospital-level Mortality Indicator (SHMI) - Deaths associated with hospitalisation 2018 https://digital.nhs.uk/data-and-information/publications/clinical-indicators/shmi. Accessed 28 December 2018.

30 Westra BL, Landman S, Yadav P, Steinbach M. Secondary Analysis of an Electronic Surveillance System Combined with Multi-focal Interventions for Early Detection of Sepsis. Appl Clin Inform. 2017;8(1):47–66.

31 Guirgis FW, Jones L, Esma R, Weiss A, McCurdy K, Ferreira J, et al. Managing sepsis: Electronic recognition, rapid response teams, and standardized care save lives. J Crit Care. 2017;40:296–302.

32 Craig P, Cooper C, Gunnell D, Haw S, Lawson K, Macintyre S et al. Using natural experiments to evaluate population health interventions: new Medical Research Council guidance. J Epidemiol Community Health 2012;66:1182–1186, 10.1136/jech-2011-200375

33 Inada-Kim M, Nsutebu E. NEWS 2: an opportunity to standardise the management of deterioration and sepsis. BMJ. 2018;360:k1260.

34 Alam N, Hobbelink EL, van Tienhoven AJ, van de Ven PM, Jansma EP, Nanayakkara PW. The impact of the use of the Early Warning Score (EWS) on patient outcomes: a systematic review Resuscitation, 85 (2014), pp. 587–594

