## Supplementary material for "Evaluating a digital sepsis alert in a London multi-site hospital network: a natural experiment using electronic health record data": who did not require antibiotics (further details in Supplementary Materials S1

**S1. Characteristics of patients who did not receive antibiotics within 24 hours of the alert**

In order to determine whether there is an association between patients suspected of sepsis given antibiotics within an appropriate time window and the status of the alert it is key to determine a sample of patients who are suspected of sepsis and for whom antibiotics are appropriate. In this study there is no agreed gold-standard for determining the patients who have sepsis, low numbers of patients have blood cultures and coding is not a reliable indicator or sepsis.

As an approximation for determining a sample of patient we are assuming that patients who need antibiotics receive them within 24 hours either side of the alert. The assumption is that patients who do not receive IV antibiotics in this time period have been seen by a clinician and there has been a clinical decision not to administer antibiotics.

In order to assess the validity of the assumption, patient characteristics and outcomes of those who did receive antibiotics were compared to those who did not receive antibiotics. These are summarised in Table 1. There are significant differences in outcomes for patients who did and did not receive antibiotics. 25% of patients who did not receive antibiotics had a long length of stay, compared to 75% of those who did. Similar differences are seen for in-hospital mortality within 30-days. A higher proportion of elderly patients received antibiotics, compared to younger patients. In addition, a higher proportion of patients with a high NEWS score received antibiotics. These significant associations suggest that more vulnerable patients are more likely to have received antibiotics. Ethnicity, deprivation and sex were not significantly associated with receipt of antibiotics, suggesting that clinical aspects of the patient and not underlying health inequalities are associated with receipt of antibiotics.

Certain patient comorbidities are associated with increased percentage receiving antibiotics. A higher percentage of dementia patient received antibiotics, although this may be confounded by age. A higher percentage of patients with metastatic cancer and myocardial infarction received antibiotics. A lower percentage of patients with a peptic ulcer received antibiotics.

A higher proportion of patients who alerted in the spring and winter received antibiotics, this may be due to a different distribution of diseases during the winter and spring months compared to summer and autumn. A lower proportion of patients admitted through the ED at SMH compared to CHX and HH. This is likely to be due to SMH providing trauma specialist care. Trauma patients are known to trigger the alert due to organ dysfunction, not related to infection. This may also be the explanation for the higher proportion of ‘suspected sepsis’ patients receiving antibiotics compared to ‘suspected **severe** sepsis’. Trauma patients may trigger **severe** sepsis alerts. This may also be the case for patients with kidney disease who have high lactate levels. The key difference between a sepsis alert and a severe sepsis alert is the presence of organ dysfunction as measured by lactate, creatinine, bilirubin and blood pressure. The associations suggest that patients not receiving antibiotics are less clinically severe.

Table S1.A Comparison of patients who alerted (either during the control (silent) or live period and did or did not receive IV antibiotics within 24 hours of the alert. p-values are derived from Chi-squared tests.

| **Factor** | **Level** | **Patients did not receive antibiotics** | | **Patients received**  **antibiotics** | | ***p-value*** |
| --- | --- | --- | --- | --- | --- | --- |
| Length of stay | <7 days | 2429 | 40.9 | 3504 | 59.1 |  |
|  | ≥7 days | 996 | 24.6 | 3059 | 75.4 | <0.0001 |
| In hospital mortality |  | 3294 | 34.9 | 6144 | 65.1 |  |
|  | Died within 30 days | 131 | 23.8 | 419 | 76.2 | <0.0001 |
| agegrpsimp18-44 |  | 740 | 40.2 | 1102 | 59.8 |  |
| agegrpsimp45-64 |  | 932 | 36.2 | 1642 | 63.8 |  |
| agegrpsimp65-69 |  | 297 | 34.7 | 560 | 65.3 |  |
| agegrpsimp70-74 |  | 333 | 32.6 | 690 | 67.4 |  |
| agegrpsimp75-79 |  | 351 | 33.4 | 700 | 66.6 |  |
| agegrpsimp80-84 |  | 312 | 30.3 | 717 | 69.7 |  |
| agegrpsimp85+ |  | 460 | 28.5 | 1152 | 71.5 | <0.0001 |
| Sex | Female | 1606 | 33.7 | 3155 | 66.3 |  |
|  | Male | 1819 | 34.8 | 3408 | 65.2 | 0.27 |
| Ethnicity | Asian | 179 | 35.2 | 329 | 64.8 |  |
|  | Black | 343 | 34.1 | 664 | 65.9 |  |
|  | NotKnown | 421 | 33.5 | 836 | 66.5 |  |
|  | Other | 696 | 35.4 | 1272 | 64.6 |  |
|  | White | 1786 | 34.0 | 3462 | 66.0 | 0.78 |
| CHF | Absent | 2968 | 34.7 | 5596 | 65.3 |  |
|  | Present | 457 | 32.1 | 967 | 67.9 | 0.06 |
| Dementia | Absent | 3160 | 35.3 | 5787 | 64.7 |  |
|  | Present | 265 | 25.5 | 776 | 74.5 | <0.0001 |
| Uncomplicated diabetes | Absent | 2581 | 33.9 | 5031 | 66.1 |  |
|  | Present | 844 | 35.5 | 1532 | 64.5 | 0.15 |
| Complicated diabetes | Absent | 3264 | 34.3 | 6257 | 65.7 |  |
|  | Present | 161 | 34.5 | 306 | 65.5 | 0.97 |
| HIV | Absent | 3389 | 34.3 | 6495 | 65.7 |  |
|  | Present | 36 | 34.6 | 68 | 65.4 | 1 |
| Liver Mild | Absent | 3187 | 34.2 | 6138 | 65.8 |  |
|  | Present | 238 | 35.9 | 425 | 64.1 | 0.39 |
| Liver Severe | Absent | 3361 | 34.4 | 6414 | 65.6 |  |
|  | Present | 64 | 30.0 | 149 | 70.0 | 0.21 |
| Metastatic cancer | Absent | 3246 | 34.8 | 6082 | 65.2 |  |
|  | Present | 179 | 27.1 | 481 | 72.9 | <0.0001 |
| MI | Absent | 3138 | 34.7 | 5914 | 65.3 |  |
|  | Present | 287 | 30.7 | 649 | 69.3 | 0.016 |
| Paralysis | Absent | 3295 | 34.3 | 6305 | 65.7 |  |
|  | Present | 130 | 33.5 | 258 | 66.5 | 0.78 |
| Peptic Ulcer | Absent | 3360 | 34.2 | 6474 | 65.8 |  |
|  | Present | 65 | 42.2 | 89 | 57.8 | 0.045 |
| Pulmonary | Absent | 2302 | 33.9 | 4481 | 66.1 |  |
|  | Present | 1123 | 35.0 | 2082 | 65.0 | 0.29 |
| PVD | Absent | 3228 | 34.5 | 6133 | 65.5 |  |
|  | Present | 197 | 31.4 | 430 | 68.6 | 0.12 |
| Renal | Absent | 3063 | 34.6 | 5793 | 65.4 |  |
|  | Present | 362 | 32.0 | 770 | 68.0 | 0.088 |
| Rheumatic | Absent | 3310 | 34.4 | 6315 | 65.6 |  |
|  | Present | 115 | 31.7 | 248 | 68.3 | 0.31 |
| Stroke | Absent | 2916 | 34.2 | 5616 | 65.8 |  |
|  | Present | 509 | 35.0 | 947 | 65.0 | 0.58 |
| Deprivation | Least deprived | 873 | 34.4 | 1663 | 65.6 |  |
|  |  | 928 | 33.3 | 1856 | 66.7 |  |
|  |  | 839 | 34.7 | 1579 | 65.3 |  |
|  |  | 392 | 34.6 | 740 | 65.4 |  |
|  | Most deprived | 164 | 33.5 | 326 | 66.5 |  |
|  | Missing | 229 | 36.5 | 399 | 63.5 | 0.73 |
| News score | Zero | 110 | 42.0 | 152 | 58.0 |  |
|  | Low | 1689 | 42.2 | 2310 | 57.8 |  |
|  | Medium | 624 | 32.3 | 1307 | 67.7 |  |
|  | High | 454 | 19.9 | 1824 | 80.1 |  |
|  | Missing | 548 | 36.1 | 970 | 63.9 | <0.0001 |
| Season | Autumn | 1189 | 37.8 | 1957 | 62.2 |  |
|  | Winter | 1074 | 31.3 | 2353 | 68.7 |  |
|  | Summer | 400 | 36.2 | 704 | 63.8 |  |
|  | Spring | 762 | 33.0 | 1549 | 67.0 | <0.0001 |
| Severity | Suspected sepsis | 1424 | 27.6 | 3744 | 72.4 |  |
|  | Suspected severe sepsis | 2001 | 41.5 | 2819 | 58.5 | <0.0001 |
| Site | A (CHX) | 1574 | 31.5 | 3420 | 68.5 |  |
|  | B (HH) | 233 | 32.5 | 485 | 67.5 |  |
|  | C (SMH) | 1618 | 37.8 | 2658 | 62.2 | <0.0001 |

**S2. Inverse probability of treatment weighting analysis**

We calculated propensity scores for patients using logistic regression to predict alert status (‘live’ (visible to clinicians) and ‘silent’ (control)). Factors included in the model to estimate propensity scores were the confounders a priori selected to be included in the main outcome analysis. Then, for each patient inpatient encounter, a weight defined as the inverse of the probability the treatment they had received was calculated. In the paper we report the marginal odds ratios (ORs) for each of the three outcomes based on weighted logistic regression.

The propensity score was first estimated with a logistic regression (multilevel with area the alert was introduced as a cluster in a random effect model for death) including all the variables listed in Table S1. For cohort A, used to model death, two propensity score models were used: ED encounters and inpatient encounters. The sample of patient encounters were different for each of the three outcomes, we therefore estimated the propensity score for each sample separately.

For the sample of patients included in the model for the outcome death, prior to propensity score weighting, deprivation quintile, most recent NEWS score, season, hospital site and renal disease were unbalanced (standardised difference >20%), additionally ethnicity, age, severity, pulmonary disease and metastatic cancer were unbalanced at a lower level (standardised difference >10%). The ‘cluster’ was highly unbalanced prior to propensity score weighting, and remained high after just above weighting. The high levels of unbalance for site and season are likely to be due to the phased introduction of the alert. All other variables are now balanced. For the samples of patients used in the models for length of stay (LOS) and timely antibiotics only season and severity of alert were unbalanced. After weighting on the propensity score all variables were balanced.

Table S2.1 .A Balance of the covariates before and after propensity score adjustment

| **Variable** | ***Standardised Mean Difference (%)*** | | | | | |
| --- | --- | --- | --- | --- | --- | --- |
|  | ***Model for death*** | | ***Model for LOS*** | | ***Model for timely antibiotics*** | |
|  | ***Before IPTW*** | ***After IPTW*** | ***Before IPTW*** | ***After IPTW*** | ***Before IPTW*** | ***After IPTW*** |
| Age group | *13.0* | 11.5 | 3.8 | 1.5 | 3.8 | 2.0 |
| Sex | 2.0 | 0.4 | 8.7 | 0.2 | 6.2 | 0.7 |
| Hospital site | **48.3** | **11.5** | 5.3 | 1.2 | 0.9 | 0.7 |
| Season | **58.8** | *6.0* | **78.2** | 3.0 | **71.9** | 2.9 |
| Ethnicity | *12.8* | 2.7 | 5.0 | 1.2 | 7.5 | 2.3 |
| Most recent NEWS score | **23.3** | *17.7* | *12.1* | 2.7 | *11.9* | 2.4 |
| Severity (suspected sepsis or suspected severe sepsis) | *10.3* | 3.1 | **21.7** | 2.6 | **22.8** | 2.2 |
| MI | 7.2 | 8.4 | 2.7 | <0.01 | 4.1 | 2.0 |
| CHF | 5.9 | 9.4 | 2.3 | 1.3 | 4.5 | 0.6 |
| PVD | 6.9 | 1.2 | 1.0 | 1.0 | 0.4 | 0.8 |
| Stroke | 0.8 | 1.7 | 0.7 | 0.6 | 3.4 | 0.5 |
| Dementia | 8.0 | 1.8 | 2.9 | 0.7 | 0.2 | 1.4 |
| Pulmonary | *16.0* | 1.5 | 2.3 | 0.7 | 2.9 | 0.8 |
| Rheumatic | 0.3 | 4.0 | 2.2 | 0.4 | 6.0 | 2.3 |
| Peptic ulcer | 1.8 | 7.3 | 1.1 | 0.3 | 0.1 | 1.2 |
| Liver disease (mild) | 2.0 | 0.6 | 0.7 | < 0.01 | 2.1 | 0.9 |
| Diabetes (uncomplicated) | 1.1 | 4.6 | 1.8 | 1.0 | 2.0 | 1.0 |
| Diabetes (complicated) | 8.8 | 11.3 | 2.9 | 0.7 | 0.8 | 0.6 |
| Paralysis | 1.9 | 4.8 | 1.8 | 0.4 | 4.2 | 0.4 |
| Renal disease | **21.6** | *14.2* | 0.6 | 0.9 | 1.6 | 0.7 |
| Liver disease (severe) | 1.9 | 0.6 | 0.5 | 0.7 | 0.2 | 1.0 |
| Metastatic cancer | *10.6* | 1.2 | 2.6 | 0.7 | 1.4 | 0.3 |
| HIV | 0.5 | 3.1 | 2.1 | 0.3 | 4.2 | 1.6 |
| Deprivation quintile | **22.4** | *9.1* | 3.1 | 1.4 | 4.6 | 2.2 |
| Cluster (ED, acute, haematology or ‘other’) | **166.1** | **20.9** |  |  |  |  |

### S3. Full results of propensity score inversely weighted logistic models

**Table S3.A Results of logistic regression of in-hospital mortality within 30 days.**

| Factor | Level | OR | CI | std.error_p | p.value_p |
| --- | --- | --- | --- | --- | --- |
| Alert Status | NA | 0.762 | (0.695, 0.836) | 0.047 | <0.0001 |
| Age group | 18-44 (reference) | NA | NA | NA | NA |
|  | 45-64 | 1.916 | (1.541, 2.382) | 0.111 | <0.0001 |
|  | 65-69 | 2.471 | (1.930, 3.165) | 0.126 | <0.0001 |
|  | 70-74 | 2.808 | (2.199, 3.585) | 0.125 | <0.0001 |
|  | 75-79 | 3.833 | (3.031, 4.848) | 0.12 | <0.0001 |
|  | 80-84 | 3.969 | (3.121, 5.046) | 0.123 | <0.0001 |
|  | 85 and over | 5.603 | (4.457, 7.044) | 0.117 | <0.0001 |
| Gender | Male (female is reference) | 1.184 | (1.079, 1.300) | 0.047 | 0.0004 |
| Ethnicity | White (reference) | NA | NA | NA | NA |
|  | Black | 0.743 | (0.625, 0.882) | 0.088 | 0.0007 |
|  | Not Known | 0.898 | (0.786, 1.026) | 0.068 | 0.115 |
|  | Other | 0.67 | (0.586, 0.766) | 0.068 | <0.0001 |
|  | Asian | 0.605 | (0.485, 0.756) | 0.113 | 0<0.0001 |
| Deprivation quintile | Least deprived quintile (reference) | NA | NA | NA | NA |
|  | NA | 0.995 | (0.872, 1.135) | 0.067 | 0.937 |
|  | NA | 1.115 | (0.977, 1.273) | 0.068 | 0.107 |
|  | NA | 0.858 | (0.729, 1.010) | 0.083 | 0.066 |
|  | Most deprived | 0.649 | (0.521, 0.810) | 0.113 | 0.0001 |
|  | Not known | 1.182 | (0.934, 1.496) | 0.12 | 0.165 |
| MI | | 1.066 | (0.929, 1.222) | 0.07 | 0.365 |
| CHF | | 2.119 | (1.896, 2.367) | 0.057 | <0.0001 |
| PVD | | 1.345 | (1.161, 1.558) | 0.075 | <0.0001 |
| Stroke | | 2.06 | (1.841, 2.305) | 0.057 | <0.0001 |
| Dementia | | 1.404 | (1.226, 1.609) | 0.069 | <0.0001 |
| Pulmonary | | 0.786 | (0.706, 0.874) | 0.055 | <0.0001 |
| Rheumatic | | 1.025 | (0.808, 1.302) | 0.122 | 0.837 |
| Peptic ulcer | | 1.09 | (0.801, 1.483) | 0.157 | 0.584 |
| Liver disease – mild | | 1.592 | (1.332, 1.903) | 0.091 | <0.0001 |
| Diabetes - uncomplicated | | 0.705 | (0.629, 0.790) | 0.058 | <0.0001 |
| Diabetes – complicated | | 0.669 | (0.539, 0.831) | 0.111 | 0.0003 |
| Paralysis | | 1.749 | (1.479, 2.070) | 0.086 | <0.0001 |
| Renal | | 1.339 | (1.186, 1.512) | 0.062 | <0.0001 |
| Liver disease – severe | | 4.723 | (3.800, 5.871) | 0.111 | <0.0001 |
| Metastatic cancer | | 6.685 | (5.909, 7.563) | 0.063 | <0.0001 |
| HIV | | 0.163 | (0.067, 0.397) | 0.454 | <0.0001 |
| Trust site - reference | A | NA | NA | NA | NA |
|  | B | 0.833 | (0.717, 0.966) | 0.076 | 0.016 |
|  | C | 0.905 | (0.807, 1.015) | 0.058 | 0.087 |
| Season of admission | Spring (reference) | NA | NA | NA | NA |
|  | Summer | 0.784 | (0.661, 0.930) | 0.087 | 0.005 |
|  | Autumn | 1.05 | (0.924, 1.193) | 0.065 | 0.453 |
|  | Winter | 1.257 | (1.110, 1.424) | 0.063 | 0.0003 |
| Severity | Suspected Sepsis (reference) | NA | NA | NA | NA |
|  | Suspected Severe Sepsis | 1.456 | (1.327, 1.596) | 0.047 | <0.0001 |
| News score | Zero (reference) | NA | NA | NA | NA |
|  | Low | 1.949 | (1.402, 2.711) | 0.168 | <0.0001 |
|  | Medium | 3.156 | (2.250, 4.426) | 0.173 | <0.0001 |
|  | High | 4.681 | (3.347, 6.545) | 0.171 | <0.0001 |
|  | Missing | 5.089 | (3.616, 7.162) | 0.174 | <0.0001 |

**Table S3.B Results of propensity score weighted logistic regression of length of stay – binary indicator of long stay patients (PILOS > 6 days).**

| **Factor** | **Level** | | **OR** | **CI** | **std.error_p** | **p.value_p** |
| --- | --- | --- | --- | --- | --- | --- |
| **Alert Status** | **SILENT IS REFERENCE** | | 0.931 | (0.876, 0.989) | 0.031 | 0.020 |
| **Age group** | **18-44 (reference)** | | NA | NA | NA | NA |
|  | **45-64** | | 1.77 | (1.595, 1.963) | 0.053 | <0.0001 |
|  | **65-69** | | 1.976 | (1.728, 2.260) | 0.068 | <0.0001 |
|  | **70-74** | | 2.053 | (1.807, 2.333) | 0.065 | <0.0001 |
|  | **75-79** | | 2.626 | (2.306, 2.990) | 0.066 | <0.0001 |
|  | **80-84** | | 2.595 | (2.273, 2.962) | 0.068 | <0.0001 |
|  | **85 and over** | | 3.33 | (2.944, 3.766) | 0.063 | <0.0001 |
| **Gender** | **Male (female is reference)** | | 1.091 | (1.025, 1.160) | 0.032 | 0.006 |
| **Ethnicity** | **White (reference)** | |  |  |  |  |
|  | **Black** | | 1.094 | (0.983, 1.217) | 0.055 | 0.010 |
|  | **Not Known** | | 1.066 | (0.969, 1.172) | 0.049 | 0.190 |
|  | **Other** | | 0.838 | (0.770, 0.912) | 0.043 | <0.0001 |
|  | **Asian** | | 0.969 | (0.840, 1.118) | 0.073 | 0.664 |
| **Deprivation quintile** | **Least deprived quintile (reference)** | |  |  |  |  |
|  |  |  | 0.996 | (0.914, 1.086) | 0.044 | 0.932 |
|  |  | | 1.121 | (1.025, 1.227) | 0.046 | 0.013 |
|  |  | | 1.211 | (1.083, 1.356) | 0.057 | 0.0008 |
|  | **Most deprived** | | 1.03 | (0.883, 1.201) | 0.079 | 0.710 |
|  | **Not known** | | 0.835 | (0.721, 0.968) | 0.075 | 0.017 |
| **MI** | | | 0.904 | (0.812, 1.007) | 0.055 | 0.065 |
| **CHF** | | | 1.583 | (1.446, 1.732) | 0.046 | <0.0001 |
| **PVD** | | | 1.384 | (1.222, 1.566) | 0.063 | <0.0001 |
| **Stroke** | | | 1.335 | (1.218, 1.465) | 0.047 | <0.0001 |
| **Dementia** | | | 1.21 | (1.088, 1.345) | 0.054 | 0.0004 |
| **Pulmonary** | | | 0.995 | (0.930, 1.065) | 0.035 | 0.886 |
| **Rheumatic** | | | 0.971 | (0.829, 1.137) | 0.081 | 0.716 |
| **Peptic ulcer** | | | 1.045 | (0.821, 1.330) | 0.123 | 0.721 |
| **Liver disease - mild** | | | 1.535 | (1.359, 1.733) | 0.062 | <0.0001 |
| **Diabetes - uncomplicated** | | | 0.969 | (0.901, 1.043) | 0.038 | 0.408 |
| **Diabetes - complicated** | | | 1.106 | (0.951, 1.287) | 0.077 | 0.190 |
| **Paralysis** | | | 1.815 | (1.545, 2.132) | 0.082 | <0.0001 |
| **Renal** | | | 1.046 | (0.945, 1.157) | 0.051 | 0.386 |
| **Liver disease - severe** | | | 3.153 | (2.562, 3.880) | 0.106 | <0.0001 |
| **Metastatic cancer** | | | 1.683 | (1.495, 1.895) | 0.061 | <0.0001 |
| **HIV** | | | 1.265 | (0.946, 1.693) | 0.149 | 0.113 |
| **Trust site - reference** | | **A (CHX)** |  |  |  |  |
| **Trust site** | | **B (HH)** | 4.134 | (3.633, 4.704) | 0.066 | <0.0001 |
| **Trust site** | | **C (SMH)** | 1.178 | (1.100, 1.263) | 0.035 | <0.0001 |
| **Season of admission** | **Spring (reference)** | |  |  |  |  |
|  | **Summer** | | 0.673 | (0.600, 0.753) | 0.058 | <0.0001 |
|  | **Autumn** | | 0.971 | (0.893, 1.055) | 0.043 | 0.483 |
|  | **Winter** | | 1.063 | (0.979, 1.154) | 0.042 | 0.148 |
| **Severity** | **Suspected Sepsis** | |  |  |  |  |
|  | **Suspected Severe Sepsis** | | 1.003 | (0.942, 1.067) | 0.032 | 0.936 |
| **News score** | **Zero (reference)** | |  |  |  |  |
|  | **Low** | | 1.207 | (0.984, 1.482) | 0.104 | 0.071 |
|  | **Medium** | | 1.322 | (1.071, 1.632) | 0.108 | 0.009 |
|  | **High** | | 1.941 | (1.575, 2.391) | 0.107 | <0.0001 |
|  | **Missing** | | 1.618 | (1.307, 2.002) | 0.109 | 0.00001 |

**Table S3.C Results of propensity score weighted logistic regression of timely antibiotics – binary indicator of IV antibiotics 12 hours before the alert or one hour after the alert.**

| **Factor** | **Level** | OR | CI | SE | p.value |
| --- | --- | --- | --- | --- | --- |
| **Alert Status** | **SILENT IS REFERENCE** | 1.203 | (1.119, 1.292) | 0.037 | <0.0001 |
| **Age group** | **18-44 (reference)** | 1.000 | NA | NA | NA |
|  | **45-64** | 1.064 | (0.923, 1.226) | 0.072 | 0.393 |
|  | **65-69** | 1.262 | (1.043, 1.527) | 0.097 | 0.017 |
|  | **70-74** | 1.106 | (0.923, 1.324) | 0.092 | 0.274 |
|  | **75-79** | 1.072 | (0.892, 1.288) | 0.094 | 0.458 |
|  | **80-84** | 1.084 | (0.898, 1.308) | 0.096 | 0.402 |
|  | **85 and over** | 1.113 | (0.936, 1.323) | 0.088 | 0.226 |
| **Gender** | **Male (female is reference)** | 0.977 | (0.893, 1.068) | 0.046 | 0.609 |
| **Ethnicity** | **White (reference)** | 1.000 | NA | NA | NA |
|  | **Black** | 1.134 | (0.972, 1.324) | 0.079 | 0.111 |
|  | **Not Known** | 1.126 | (0.983, 1.289) | 0.069 | 0.087 |
|  | **Other** | 0.97 | (0.859, 1.095) | 0.062 | 0.619 |
|  | **Asian** | 1.066 | (0.864, 1.315) | 0.107 | 0.550 |
| **Deprivation quintile** | **Least deprived quintile (reference)** | 1.000 | NA | NA | NA |
| **quintile** |  | 0.984 | (0.870, 1.113) | 0.063 | 0.800 |
|  |  | 0.743 | (0.651, 0.849) | 0.067 | <0.0001 |
|  |  | 0.835 | (0.708, 0.985) | 0.084 | 0.032 |
|  | **Most deprived** | 0.647 | (0.515, 0.813) | 0.116 | 0.00018 |
|  | **Not known** | 1.000 | (0.813, 1.230) | 0.106 | 0.998 |
| **MI** | | 0.902 | (0.768, 1.059) | 0.082 | 0.207 |
| **CHF** | | 1.077 | (0.940, 1.234) | 0.069 | 0.283 |
| **PVD** | | 1.02 | (0.849, 1.227) | 0.094 | 0.830 |
| **Stroke** | | 0.94 | (0.815, 1.086) | 0.073 | 0.40116 |
| **Dementia** | | 1.261 | (1.080, 1.473) | 0.079 | 0.003 |
| **Pulmonary** | | 1.003 | (0.908, 1.107) | 0.051 | 0.955 |
| **Rheumatic** | | 0.866 | (0.693, 1.083) | 0.114 | 0.207 |
| **Peptic ulcer** | | 1.047 | (0.689, 1.589) | 0.213 | 0.831 |
| **Liver disease - mild** | | 0.618 | (0.509, 0.749) | 0.098 | <0.0001 |
| **Diabetes - uncomplicated** | | 0.953 | (0.855, 1.062) | 0.055 | 0.381 |
| **Diabetes - complicated** | | 0.74 | (0.587, 0.932) | 0.118 | 0.011 |
| **Paralysis** | | 1.215 | (0.952, 1.549) | 0.124 | 0.117 |
| **Renal** | | 1.103 | (0.949, 1.283) | 0.077 | 0.202 |
| **Liver disease - severe** | | 0.49 | (0.351, 0.685) | 0.171 | <0.0001 |
| **Metastatic cancer** | | 0.952 | (0.802, 1.131) | 0.088 | 0.578 |
| **HIV** | | 0.377 | (0.236, 0.603) | 0.24 | <0.0001 |
| **Trust site** | **A** | 1.000 | NA | NA | NA |
|  | **B** | 0.71 | (0.592, 0.850) | 0.092 | 0.0002 |
|  | **C** | 0.588 | (0.531, 0.651) | 0.052 | <0.0001 |
| **Season of admission** | **Spring (reference)** | 1.000 | NA | NA | NA |
|  | **Summer** | 2.006 | (1.716, 2.346) | 0.08 | <0.0001 |
|  | **Autumn** | 1.012 | (0.897, 1.142) | 0.062 | 0.846 |
|  | **Winter** | 0.903 | (0.803, 1.015) | 0.06 | 0.088 |
| **Severity** | **Suspected Sepsis (reference)** | 1.000 | NA | NA | NA |
|  | **Suspected Severe Sepsis** | 0.714 | (0.651, 0.783) | 0.047 | <0.0001 |
| **News score** | **Zero (reference)** | 1.000 | NA | NA | NA |
|  | **Low** | 1.054 | (0.757, 1.469) | 0.169 | 0.754 |
|  | **Medium** | 1.879 | (1.341, 2.632) | 0.172 | 0.0002 |
|  | **High** | 2.922 | (2.090, 4.086) | 0.171 | <0.0001 |
|  | **Missing** | 1.566 | (1.114, 2.202) | 0.174 | 0.010 |
